# Evaluating Genetic Regulators of MicroRNAs Using Machine Learning Models

**DOI:** 10.1101/2025.01.09.632215

**Authors:** Mert Cihan, Uchenna Alex Anyaegbunam, Steffen Albrecht, Miguel A Andrade-Navarro, Maximilian Sprang

## Abstract

This study explores the genetic regulators of microRNAs (miRNAs) using an ensemble of machine learning models to predict miRNA expression levels from gene expression data. Employing ridge regression, we accurately predicted the expression of 353 human miRNAs (R^2^ > 0.5), revealing robust miRNA-gene regulatory relationships. By analyzing the coefficients of these predictive models, we identified genetic regulators for each miRNA and highlighted the multifactorial nature of miRNA regulation.

Further network analysis uncovered that miRNAs with higher predictive accuracy are more densely connected to their top predictive genes, reflecting strong regulatory control within miRNA-gene networks. To refine these insights, we filtered the gene-miRNA interaction networks to identify miRNAs specifically associated with enriched pathways, such as synaptic function and cardiovascular processes. From this pathway-centric analysis, we present a curated list of miRNAs and their genetic regulators, pinpointing their activity within distinct biological contexts.

Additionally, our study provides a comprehensive set of metrics and coefficients for the genes most predictive of miRNA expression, along with a filtered subnetwork of miRNAs linked to specific pathways and phenotypes. By integrating miRNA expression predictors with network analysis and pathway enrichment, this work advances our understanding of miRNA regulatory mechanisms and their roles across distinct biological systems.

## Introduction

MicroRNAs (miRNAs) play a critical role in the regulation of gene expression by binding to target messenger RNAs (mRNAs) and either promoting their degradation or inhibiting their translation [1]. These small, non-coding RNAs are involved in a wide array of cellular processes, including development, differentiation, and apoptosis, making them essential for maintaining cellular homeostasis [2,3].

Accurate profiling of miRNA expression is crucial for understanding miRNA functions. To predict miRNA targets, among other methods, the negative correlation between miRNA and mRNA expression is used to identify potential novel miRNA target binding sites on genes [4–6]. By mapping these interactions, researchers can elucidate how miRNAs influence various cellular pathways and processes, highlighting their potential as therapeutic targets and biomarkers [7,8]. This characterization not only relies on direct binding site identification but also on integration with annotation databases, high-throughput experimental validation, evolutionary conservation studies, and network-based analysis [3,9–12].

For the quantification of miRNA expression, multiple experimental methods can be used, each facing its own challenges.

Most current small RNA-seq protocols rely on adapter ligation and PCR amplification for library construction, which introduces biases in miRNA representation, affecting the accuracy and reliability of miRNA profiling [13–15]. Additionally, the vast range of sequencing reads produced can lead to issues such as false high fold changes from very small expression values and errors stemming from de-multiplexing and alignment ambiguities when compared to quantitative RT-PCR, which typically offers higher precision [16].

Moreover, integrating miRNA expression data from various platforms, such as microarrays, next-generation sequencing, bead-based detection systems, single-molecule measurements, and quantitative RT-PCR, often results in inconsistencies due to the distinct biases and error profiles inherent in each method. A major obstacle affecting the reliability of miRNA expression data is the lack of validated reference controls for data normalization, which leads to significant variability and hinders cross-study comparisons [13,17–19].

In this study, we use gene expression data from RNA sequencing to predict miRNA expression levels, offering a novel approach that leverages the correlations between gene and miRNA expressions to build predictive machine learning models, providing a more accessible and accurate computational alternative to direct miRNA measurement. By doing so, it allows us to infer miRNA activity and its regulatory impact on genes, facilitating deeper insights into both cellular mechanisms and disease pathways.

To this end, we applied ridge regression [20], a regularization technique suited for handling multicollinearity and high-dimensional data, to predict miRNA levels from RNA-seq data obtained from The Cancer Genome Atlas (TCGA) [21] from both normal and cancer tissues. By evaluating the model’s performance using metrics such as R-squared (R^2^) values, we were able to assess the accuracy and reliability of our predictions. This approach not only demonstrated the feasibility of predicting miRNA expression from gene data but also provided insights into the complex regulatory relationships between genes and miRNAs. By analyzing the regression coefficients, we identified predictive genes for each miRNA considered, revealing key regulatory elements within the gene-miRNA network. Subsequent network analysis, incorporating miRNA binding data, enabled us to map out intricate interactions and pathways to characterize the functional relevance and biological implications of these predictive genes.

## Methods

### Data Collection

We sourced expression data from TCGA [21], selecting samples that included both miRNA sequencing and RNA-seq data. Our dataset comprised 10,464 matched expression profiles, with 9,828 derived from tumor tissues and 636 from normal tissues. To ensure uniformity and comparability across all samples, we used normalized expression values using Transcripts Per Million (TPMs). This normalization method accounts for differences in sequencing depth and gene length, providing a standardized framework for subsequent analysis.

### miRNA Expression Modeling and Data Preparation

To model miRNA expression levels, we utilized gene expression values as predictive features. The initial data preparation phase involved several key steps: loading the RNA-seq and miRNA-seq datasets and filtering the feature matrix by removing rows with more than 1,050 zeros and 1,050 NaNs, a threshold set to 10% of the total samples. For the label matrix, we applied a less stringent threshold of 10,000 zeros and 10,000 NaNs. Given that miRNA expression is often tissue-specific and generally lower in abundance, this threshold allows us to retain miRNAs present in at least 325 samples, capturing approximately 70% of miRNAs, even those expressed at lower levels. This filtering strategy enabled elimination of non-informative features and labels, enhancing the robustness of subsequent analyses. We also removed features with zero variance to focus on informative predictors.

Next, we applied a Z-transformation using the *StandardScaler* function from the scikit-learn library [22] to standardize the features so that the values of different features are on the same scale to ensure that the regularization applies equally to all coefficients, which is important for the performance of the ridge regression. Each miRNA’s dataset was split into training (80%) and test (20%) subsets, and ridge regression was applied using the *Ridge* function. This regularization technique is effective in managing multicollinearity and high-dimensional datasets, chosen for its ability to prevent overfitting and enhance prediction stability. The regularization parameter (alpha) was set at 11,000 after initial tuning after an initial grid search with 5-fold cross-validation on the same training set over a range of values (1, 10, 100, 1000, 10,000, 11,000, and 15,000), selecting 11,000 as it yielded the highest number of miRNAs with R^2^ values greater than 0.5 while maintaining the lowest mean absolute error. This methodology ensures that the test set remained completely independent throughout the training and model optimization process to provide an unbiased evaluation of model performance.

Interestingly, running the same model with the top 3% of features with the highest absolute coefficients yielded similar results (Supplementary Tables S1 and S2). While we focused on these top features for downstream analysis, we retained all features for modeling, as computing efficiency was not significantly impacted, and the overall results improved.

### Model Performance Evaluation

The models were trained on the training set and evaluated on the test set using metrics such as R^2^, median absolute error, and mean absolute error (Supplementary Tables S1 and S2). These metrics provided a comprehensive view of model performance across different miRNAs. The final dataset consisted of 21,044 features (mRNA expression data) and 1,300 miRNAs as targets. All features were retained in the final models, as feature exclusion did not improve model efficiency, and retaining them enhanced predictive accuracy. The top 3% genes with the highest absolute coefficients for miRNAs predicted with R^2^ >0.5 are provided in Supplementary Table S3. The entire modeling process was conducted using Python and R.

### Network construction and subsetting

Following the predictive modeling, we constructed a network of miRNAs and their target genes. This network was built by combining experimentally validated interactions from the miRTarBase [23], DIANA-TarBase [24] and TRANSFAC databases [25]. Additionally, we complemented these interactions with predicted conserved interactions from the TargetScanHuman 8.0 [9] with conservation scores higher than 0.5 to ensure a comprehensive representation of miRNA-gene interactions. In this network, each node represents a gene or miRNA, and the edges represent the interactions between them.

### Connectivity Metrics

To analyze miRNA connectivity within the network, we used the biomaRt package [26] to classify gene biotypes as protein-coding or long non-coding RNAs (lncRNAs). Key connectivity metrics were computed using the igraph package [27]: *degree* assessed the number of direct interactions, *betweenness* measured the role of miRNAs as bridges in the network, and *evcent* evaluated their influence based on connections. Communities were computed with *cluster_louvain*.

Pearson correlation coefficients between R^2^ values and centrality measures were calculated using the *cor* function. Network metrics for miRNA-gene regulatory interactions, including connectivity measures, centrality scores, and community structure, are provided in Supplementary Tables S4 and S5.

### Gene Ontology (GO) Term Analysis

For the Gene Ontology (GO) term analysis, we focused on the top 3% of the most predictive genes for each miRNA, selected based on R^2^ values greater than 0.5. This analysis concentrated on significant findings with adjusted p-values smaller than 0.05. Utilizing the *enrichGO* function from clusterProfiler [28], we examined Biological Processes (BP), Molecular Functions (MF), and Cellular Components (CC) to uncover the biological and molecular underpinnings influenced by these highly predictive genes. Detailed GO term enrichment results are provided in Supplementary Table S6.

### Pathway Analysis

For the pathway analysis, we again focused on the top 3% most predictive genes for each miRNA with R^2^ values greater than 0.5. We performed enrichment analysis using the KEGG database [29], applying a threshold of adjusted p-values smaller than 0.05 to identify significant pathways (see Supplementary Tables S7 and S8). This involved utilizing the *enrichKEGG* function from the clusterProfiler [28] package to map the predictive genes to their associated biological pathways, providing insights into the regulatory frameworks governing miRNA-mediated gene expression.

### Disease Enrichment

The disease enrichment analysis was conducted using the top 3% of predictive genes for each miRNA, identified based on R^2^ values greater than 0.5. These genes were cross-referenced with the Human Phenotype Ontology database [30] through the g:Profiler web platform [31], with a p-value cutoff of 0.01 (see Supplementry Table S9).

### Organ-specific miRNA expression

For the comparison of organ-specific miRNA expression levels between miRNAs associated with signal transduction pathways and all other miRNAs with R^2^ values greater than 0.5, we utilized the TAM 2.0 database [32] to extract miRNA levels for each organ. The expression levels of the pathway-specific miRNAs were contrasted with the remaining miRNAs, and the significance of the differences was assessed using a t-test using *t*.*test* function of the R stats library.

## Results

In this study, we applied ridge regression to develop an ensemble of models to predict miRNA expression levels from RNA sequencing data. By leveraging the correlations between gene and miRNA expressions, our approach provides a computational alternative to direct miRNA measurement. Additionally, we constructed a network of miRNAs and their target genes, integrating experimentally validated interactions and predicted conserved interactions to understand the functional relevance and biological implications of these regulatory relationships better. We then used the feature coefficients from these models to identify key predictive genes, allowing us to explore the regulatory elements within the gene-miRNA network. This analysis offers a deeper insight into miRNA-gene interactions and their roles in cellular mechanisms and disease pathways.

### Model Development and Performance Assessment

We thoroughly evaluated the performance of the ridge regression models that we utilized to predict miRNA expression levels from RNA sequencing data, using various statistical metrics. The distribution of R^2^ values across all miRNAs (Figure 1A) reveals a wide range of predictive accuracies. The cumulative distribution function (CDF) overlay shows that 353 out of the 1,300 miRNAs analyzed achieve R^2^ values greater than 0.5, demonstrating strong predictive capabilities for these specific targets. This may stem from their inherently high expression levels, as reflected by the median TPM values extracted from respective TCGA samples (R^2^ ≤ 0.5: 0.14; R^2^ > 0.5: 39 ;See Methods for details).

**Figure 1:**
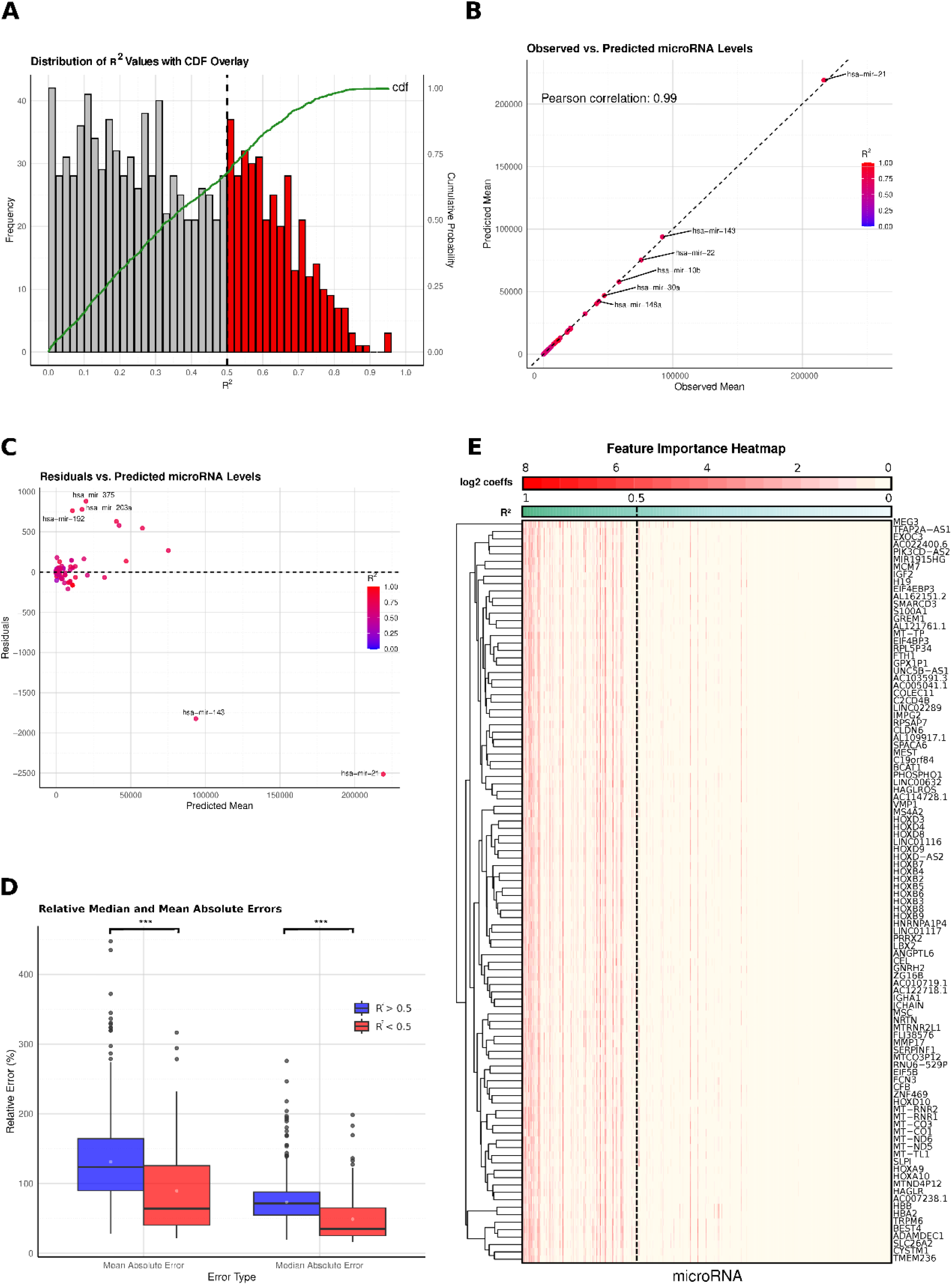
miRNA Prediction Performance and Feature Importance Analysis. (A) Distribution of R^2^ values across predicted miRNAs, with a histogram illustrating the range of prediction accuracies and cumulative probability overlayed in green. (B) Comparison of observed versus predicted mean miRNA expression levels, shown in a scatterplot with data points colored according to R^2^ values, highlighting prediction accuracy across samples. (C) Residual analysis displaying the differences between observed and predicted mean values plotted against predicted mean miRNA levels, with outliers highlighted. (D) Boxplots comparing relative errors (median and mean absolute errors) for miRNAs grouped by predicted R^2^ values (<0.5 and >0.5), providing insight into prediction reliability across different accuracy levels. (E) Clustered heatmap of the top 100 genes with the highest absolute coefficients, showing feature importance by miRNA. Genes are sorted in descending order of R^2^ values, visualizing the predictive contributions across miRNAs.

The comparison between observed and predicted miRNA levels (Figure 1B) shows a strong linear correlation, as evidenced by a Pearson correlation coefficient of 0.99, indicating the model’s proficiency in accurately capturing the mean expression levels for the majority of miRNAs, reinforcing the validity of using gene expression data as a reliable surrogate for direct miRNA measurement.

To evaluate the model’s consistency, we examined the correlation between the coefficients of variation (CV) for observed and predicted miRNA levels. Overall, this correlation was moderate (r = 0.32), indicating some alignment between observed and predicted variability. For miRNAs with R^2^ values above 0.5, the correlation was much stronger (r = 0.98), demonstrating high stability and consistency in predictions. Conversely, for miRNAs with R^2^ values < 0.5, the correlation dropped to 0.20, highlighting greater challenges in accurately predicting these miRNAs. These results reinforce the model’s robustness, especially for miRNAs with higher predictive accuracy.

Residual analysis (Figure 1C) provides additional insights into the model’s robustness and areas for improvement. While the residuals generally cluster around zero, indicating unbiased predictions across most expression levels, certain miRNAs, such as hsa-mir-21, exhibit significant deviations from the trend. These deviations are primarily associated with miRNAs that have very high expression levels, suggesting that outlier values or extreme expression levels may introduce some noise or variability into the predictive model.

The relative errors, both mean absolute error (MAE) and median absolute error (MedAE), were significantly lower for miRNAs with R^2^ > 0.5 compared to those with R^2^ < 0.5, as shown in Figure 1D, with a p-value less than 0.01. This highlights the improved predictive accuracy for miRNAs with higher R^2^ values, pointing at the model being more robust for targets with generally higher expression values (Figure 1D).

For the top 100 genes with the highest variability, the feature importance heatmap (Figure 1E) illustrates the absolute log2 values of the coefficients for miRNAs with R^2^ values greater than 0.5, revealing that many features have high coefficients. This indicates that the prediction of miRNA expression is not driven by a single gene but rather by a complex interaction among multiple genes. The presence of several genes with substantial coefficients underscores the multifactorial nature of miRNA regulation, validating the model’s strategy of using a diverse set of gene expression data to enhance predictive accuracy. Conversely, for miRNAs with R^2^ values < 0.5, there are no significantly high coefficients, suggesting a lack of strong predictive features and highlighting the challenges in predicting these miRNAs accurately (Figure 1E).

Overall, the results demonstrate that our ridge regression models provide a robust framework for predicting miRNA expression from RNA-seq data, particularly for miRNAs with clear expression patterns.

### miRNA-Gene Network Connectivity and Centrality Analysis

We analyzed the connectivity of the top 3% (632) of predictive genes, determined by the highest absolute coefficients for each miRNA, within the gene-miRNA network (Figure 2A). For miRNAs with R^2^ > 0.5, a higher proportion of predictive genes were found to be directly interacting with the miRNA (1-node distance), averaging 125 genes, compared to 102 genes for miRNAs with R^2^ < 0.5. Additionally, when examining the 3-node distance (3 degrees of separation), the difference between the two groups becomes more pronounced, with miRNAs that are better predicted (R^2^ > 0.5) showing an average of 401 connected genes, compared to 358 for those with R^2^ < 0.5. This suggests that miRNAs with higher predictive power tend to form stronger direct regulatory relationships with their target genes, highlighting the connection between prediction accuracy and regulatory interactions.

**Figure 2:**
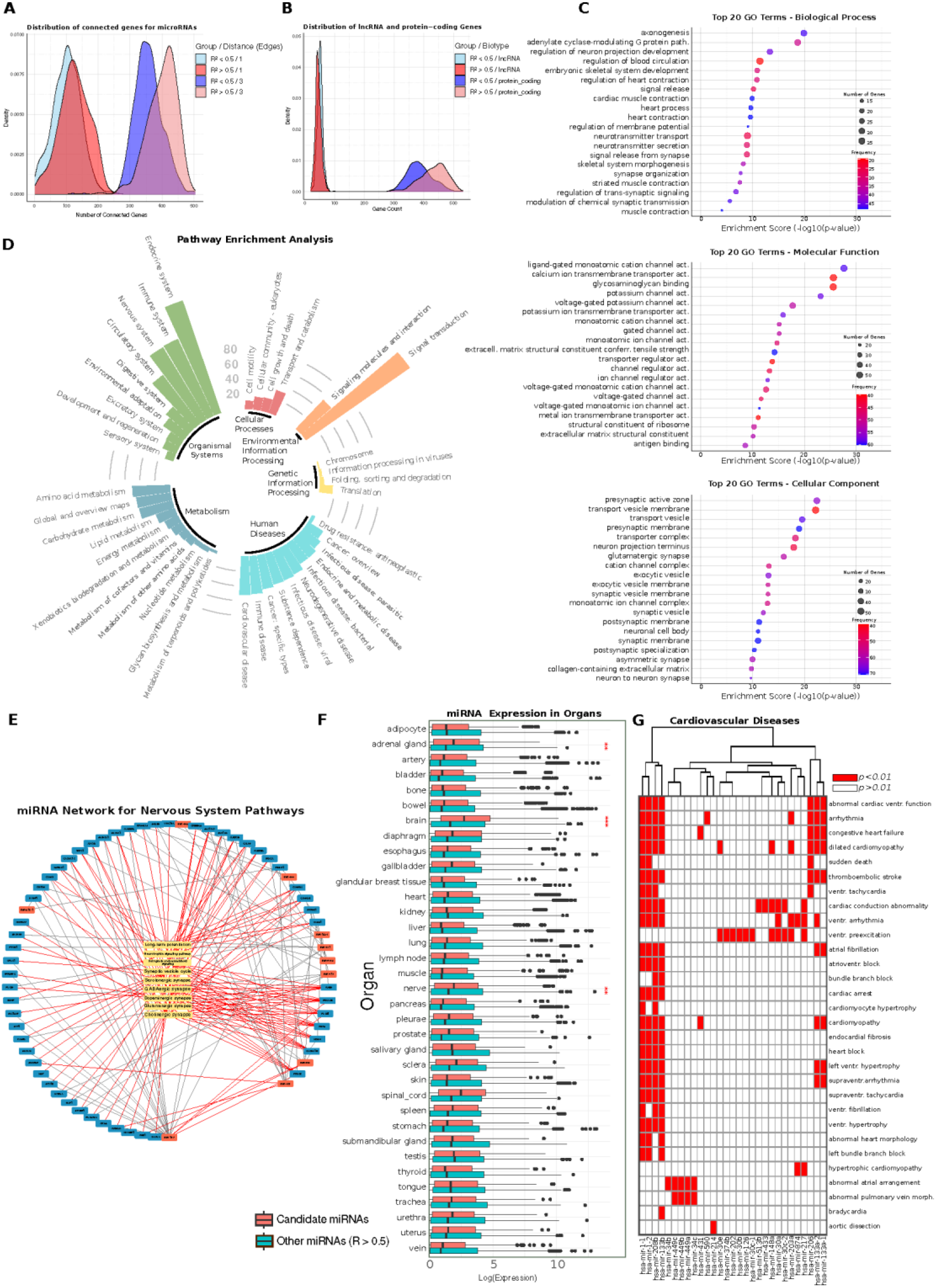
Connectivity and Functional Characterization of Predictive Genes in miRNA Networks. (A) Distribution of direct and 3-node distance gene interactions among the top 632 predictive genes for each miRNA, divided into 6. Distribution plot highlights differences in connectivity for high and low accuracy miRNAs. (B) Density distribution of gene biotypes, with long non-coding RNAs and protein-coding genes represented among the top predictive genes for each miRNA, split by R^2^ category (R^2^ > 0.5 and R^2^ < 0.5). (C) Gene Ontology (GO) term analysis results for the top 20 enriched terms in Biological Process (BP), Cellular Component (CC), and Molecular Function (MF) categories. Enrichment is based on the frequency of significance across miRNAs, using the top 632 predictive genes per miRNA (see Supplementary Table S6). (D) KEGG-pathway enrichment analysis for the top predictive genes across miRNAs, illustrated in a bar plot where the height of each bar represents the number of miRNAs significantly enriched in each pathway. This visualization highlights the pathways most frequently associated with genes that predict miRNA expression. (E) Filtered network of miRNA-gene interactions, focused on pathways related to the nervous system. Network visualizes specific interactions in nervous system-associated pathways. (F) Bar plot displaying tissue-specific expression levels of miRNAs that are filtered for signal transduction pathways and for the background of all other miRNAs predicted with R^2^ > 0.5. (G) Heatmap showing miRNAs significantly enriched in cardiovascular disease pathways (p<0.01).

We also examined the distribution of lncRNAs and protein-coding genes among the top predictive genes. Both groups, miRNAs with R^2^ > 0.5 and R^2^ < 0.5, had a similarly small proportion of lncRNAs among the top predictive genes. However, the proportion of protein-coding genes was higher for miRNAs with R^2^ > 0.5 (Figure 2B).

Analysis of the network’s communities (groups of densely connected nodes, see Methods for details) shows variability in the proportion of well-predicted miRNAs (R^2^ > 0.5) across different communities, with some communities having a higher concentration of accurately predicted miRNAs. While this observation suggests differences in the predictive relationships within these communities, no consistent pattern was observed regarding community size (nodes/edges) and prediction quality (see Supplementary Table S5).

We analyzed the relationship between miRNA expression variability and their connectivity within the network by focusing on the 55 miRNAs with a high coefficient of variation (CV > 10). Correlating their R^2^ values with different network centrality measures revealed notable relationships: a Pearson correlation of 0.49 for both degree centrality and betweenness centrality and 0.47 for eigenvector centrality. These positive correlations suggest that miRNAs with higher variability in expression tend to be predicted better when they occupy more central and influential positions in the network.

This finding implies that miRNAs with significant network connectivity—either by having numerous direct interactions (degree centrality), being central to communication pathways (betweenness centrality), or influencing other highly connected nodes (eigenvector centrality)—are more likely to exhibit predictable expression patterns. This could indicate that miRNAs deeply embedded in the regulatory network play crucial roles in maintaining network stability, which could explain why their expression is better captured by predictive models.

### Functional Enrichment of Genetic Regulators

In the GO term analysis of the top predictive genes for miRNAs with R^2^ > 0.5, we found that many terms in the biological process category are related to synaptic function and cardiovascular processes. Notably, terms such as modulation of chemical synaptic transmission, synapse organization, neurotransmitter secretion, and regulation of neuron projection development had the highest number of predictive genes associated with them. In addition, cardiovascular-related terms like cardiac muscle contraction, heart process, and regulation of blood circulation were also highly enriched, underscoring the involvement of these genes in critical physiological pathways (Figure 2C).

In the molecular function category, many of the enriched terms pertained to ion channel activity, particularly those involved in synaptic signaling. Higher-level categories like voltage-gated ion channel activity, monoatomic cation channel activity, and potassium channel activity dominated, with a large number of genes contributing to these functions. This suggests that most well-predicted miRNA families have predictive genes involved in regulating ion transport and signaling, further emphasizing their role in synaptic function and neuronal regulation (Figure 2C).

The cellular component category also reflected a strong focus on synaptic structures, with terms such as synaptic vesicle membrane, postsynaptic membrane, and neuronal cell body being the most enriched. These terms highlight the cellular environments where the predictive genes are most active, particularly in synapse-related functions. The enrichment in these synaptic components suggests that the genes associated with better-predicted miRNAs are often localized to critical regions involved in neural communication (Figure 2C).

These findings illustrate that the majority of well-predicted miRNA families have predictive genes that are heavily involved in synaptic and cardiovascular processes, as reflected by their enrichment in both functional and structural terms across the GO categories.

### Pathway enrichment

We subsequently performed pathway enrichment analysis using the KEGG database for the same set of predictive genes. Pathways significantly enriched across the majority of miRNAs include signal transduction, which involved 170 out of the 353 miRNAs considered (48%), the endocrine system with 160 miRNAs (45%), the nervous system with 113 miRNAs (32%), and cardiovascular diseases with 52 miRNAs (15%). These findings align with the GO term enrichment results, emphasizing synaptic and cardiovascular processes (Figure 2D).

We then filtered the miRNA-gene network to focus specifically on the genes associated with the pathways identified in the previous enrichment analysis, retaining only miRNAs directly connected to these genes. Additionally, we incorporated specific pathways corresponding to the enriched terms. For the nervous system, we presented this filtered network in Figure 2E, where key miRNAs such as miR-137 and miR-488 emerged as highly connected nodes within the network. This strategy resulted in the selection of 11 miRNAs, revealing a clear concentration of regulatory interactions within neural-associated pathways.

We applied the same filtering strategy to isolate pathways associated with signal transduction, which also led to the selection of 43 miRNAs. For these miRNAs, we further analyzed their tissue-specific expression patterns using the TAM 2.0 database [32]. We contrasted their expression levels with the expression profiles of all other miRNAs with R^2^ > 0.5. Through this comparison, we identified significantly higher normalized expression levels of these signal transduction - associated miRNAs in several key tissues, particularly the brain, nerve, and adrenal gland (Figure 2F).

These findings are consistent with the biological roles of signal transduction and nervous system pathways and provide additional validation of our network filtering methodology. The enriched expression in neural and endocrine-related tissues supports the functional relevance of the extracted miRNAs and highlights their potential regulatory impact within these critical systems. This underscores the biological coherence of our approach, linking the predictive genes and pathways to specific tissue contexts, thus reinforcing the importance of these miRNAs in neural and signal transduction-related processes.

### Cardiovascular Disease Associations in Predictive Gene Networks

We next conducted disease enrichment analysis for the predictive genes of miRNAs with R^2^ > 0.5, focusing on terms related to cardiovascular diseases, which were among the most prevalent in the pathway enrichment results (Figure 2G). Notably, the terms arrhythmia, abnormal cardiac ventricular function, and cardiac conduction abnormality were among the most frequent. Specifically, we observed significant enrichment of cardiovascular disease-related terms for the predictive genes of miR-1-1, miR-1-2, miR-208b, and miR-133b, indicating their strong association with various cardiovascular conditions. miR-208b, and miR-133b also appeared consistently when constructing the subnetwork for cardiovascular diseases, which included a total of 16 miRNAs. This reinforces the role of these specific miRNAs in cardiovascular regulation and highlights their potential importance in disease-associated regulatory networks.

## Discussion

In this study, we employed ridge regression to predict miRNA expression from RNA sequencing data, leveraging its strengths in handling high-dimensional data and capturing multicollinearity. Ridge regression has been widely used in genetic studies to address the challenges posed by complex datasets, particularly those involving gene regulatory networks [33–36]. Its ability to manage large numbers of correlated features while maintaining robust predictions makes it ideal for exploring the regulatory interactions between miRNAs and their target genes, which are often characterized by overlapping regulatory roles and multicollinear gene expression profiles [11,23,24].

Our ensemble of models performed well for a subset of miRNAs, with over 353 miRNAs achieving R^2^ values greater than 0.5. These miRNAs tend to have higher expression, which likely contributed to the model’s ability to capture their regulatory relationships accurately. The robustness of our model was further supported by the strong correlation between observed and predicted miRNA levels. The consistency in both the predicted mean and CV underscores the model’s ability to accurately reflect the variability and distribution of miRNA expression. Furthermore, the low residual errors, particularly for miRNAs with high expression, indicate that the model reliably predicts expression levels for these miRNAs. Among the top 100 miRNAs with the highest mean expression, only 8 were predicted with R^2^ values below 0.5, reinforcing the strength of the model in capturing the regulatory dynamics of highly expressed miRNAs.

In contrast, miRNAs with lower expression levels posed a greater challenge. This is likely due to their lower signal-to-noise ratios, making it difficult for the model to distinguish true signals from background noise. Additionally, data sparsity for these miRNAs limits the model’s ability to learn strong patterns, while higher expression miRNAs offer richer data. Lastly, regularization in ridge regression tends to shrink the coefficients of low-expression miRNAs, reducing their predictive accuracy. However, this trade-off is crucial for preventing overfitting, as the model must balance capturing meaningful patterns without allowing noise to dominate the predictions [37–39].

A critical factor in the model’s success was carefully selecting the regularization parameter (alpha), which we set to 11,000. This relatively high regularization was essential due to the inclusion of a large number of gene features. By applying this level of regularization, we were able to manage the complexity of the dataset, ensuring that the model did not overfit while maintaining stability in its predictions. The choice to retain a large number of gene features was deliberate, as it allowed the model to capture the nuanced regulatory relationships between genes and miRNAs, which are often distributed across a wide array of interactions.

While existing methods predominantly address miRNA expression imputation through techniques such as constrained least squares and GO-based similarity measures, our approach broadens the application to both healthy and tumor tissues, enhancing predictive performance without the need for imputation strategies [40,41]. By predicting miRNA expression directly from RNA-seq-derived expression matrices, our ensemble of models offers broader applicability without the need for pre-existing miRNA profiles or imputation, making it a versatile tool across different tissue types and conditions.

Our analysis of connectivity within the miRNA-gene network reveals significant relationships between prediction accuracy (R^2^ > 0.5) and the degree of direct and indirect gene interactions. miRNAs with higher predictive accuracy had more predictive genes directly connected and even more genes within 3-node distances. This strengthens the argument that miRNAs with better predictive performance tend to form stronger regulatory relationships, which are crucial for maintaining cellular homeostasis. Such relationships point to a biological coherence where the accurate prediction of miRNA expression correlates with robust gene interactions. Our network analysis, particularly the correlation between miRNA R^2^ values and centrality measures, further supports the idea that highly connected miRNAs are more predictable. Positive correlations with degree centrality and betweenness centrality suggest that miRNAs with greater regulatory influence tend to exhibit more stable and predictable expression patterns. This highlights the importance of considering miRNAs not in isolation but within the context of their broader network interactions. miRNAs with high centrality likely act as regulatory hubs, influencing a wide range of target genes across critical biological pathways.

We observed a significant enrichment of biological processes related to synaptic function and cardiovascular systems. Terms such as “synapse organization” and “cardiac muscle contraction” consistently appeared in the GO analysis for miRNAs with high R^2^ values, indicating a role in crucial physiological pathways. This is further validated by the pathway enrichment results, where signal transduction and cardiovascular pathways dominated. Filtered miRNAs, such as miR-1-1, miR-208b, and miR-133b, which showed strong associations with cardiovascular disease-related pathways, have been extensively documented as key players in their role in cardiovascular disease progression and biomarkers in literature [42–45], providing further validation of our findings and highlighting their critical role in cardiovascular regulation. miRNAs may play an essential role in the heart’s adaptability to varying physiological stimuli, allowing for rapid regulatory responses critical in maintaining cardiac rhythm and contractility [46,47]. This dynamic functionality of miRNAs might supplement transcription factors by enabling fine-tuned, quick adjustments in response to environmental or metabolic changes influencing gene expression.

In neurons, miRNAs may function as highly localized regulatory elements, helping to control mRNA pools in distant cellular locations like dendrites. This setup supports rapid, synapse-specific protein synthesis and is well-suited to the nervous system’s dynamic demands, where miRNAs act as localized regulators that inhibit protein production from stored mRNAs [48,49]. This regulatory mechanism is consistent with the presence of multiple polyadenylation sites in neural transcripts [50], allowing for flexible transcript pools finely regulated by miRNAs.

By subsetting these miRNA-gene networks based on genes predictive for miRNAs and enriched in signal transduction pathways, we identified miRNAs specifically linked to these pathways. When cross-referenced with independent databases, these selected miRNAs also showed higher expression in their respective tissues that are nerve, adrenal gland and brain.

This organ-specific expression additionally validates our methodological approach, indicating that the miRNAs identified as key players especially in synaptic and cardiovascular pathways are biologically relevant. This finding is consistent with existing research that demonstrates the tissue-specific roles of miRNAs in both the heart and brain [51,52].

One limitation of this study is the model’s reduced accuracy in predicting miRNAs with low expression levels, likely due to weaker signal-to-noise ratios and sparse data for these miRNAs. Additionally, the miRNA-gene network used in our analysis is based on current datasets, which are continuously evolving as new experimental data becomes available. As a result, some regulatory interactions may be missing, and the network may not fully capture all relevant biological relationships, potentially limiting the scope and applicability of our findings.

This study illustrates the effectiveness of ridge regression for predicting miRNA expression levels from gene expression data, offering a computational alternative to direct miRNA measurement. By analyzing the predictive gene-miRNA relationships, we revealed key insights into the functional roles of miRNAs, particularly in synaptic and cardiovascular processes. Our findings provide a foundation for further exploration of miRNA regulation in disease contexts, and the framework developed here has the potential for broader applications in miRNA-related research.

## Acknowledgements

This research did not receive any specific grant from funding agencies in the public, commercial, or not-for-profit sectors. All research activities were conducted using the resources available at Johannes Gutenberg University Mainz.

## Data Availability

The dataset supporting the conclusions of this article is included within the article and its additional supplementary tables. Supplementary Table 1 includes comprehensive benchmarking results for all features and the top 3% predictive features, a list of genetic predictors of microRNAs, network metrics, community membership of microRNAs within the network, results of gene ontology enrichment, KEGG pathway enrichment analysis, identified microRNA pathways, and disease association findings.

## Supplementary Data

Supplementary Data are available as xlsx.

